# Chi3l1 is a modulator of glioma stem cell states and a therapeutic vulnerability for glioblastoma

**DOI:** 10.1101/2021.10.01.462786

**Authors:** Charlotte Guetta-Terrier, David Karambizi, Bedia Akosman, Jia-Shu Chen, Suchitra Kamle, Eduardo Fajardo, Andras Fiser, Ritambhara Singh, Steven A. Toms, Chun Geun Lee, Jack A. Elias, Nikos Tapinos

## Abstract

Chi3l1 (Chitinase 3-like 1) is a secreted protein highly expressed in glioblastoma. Here, we show that exposure of glioma stem cells (GSCs) to Chi3l1 reduces the CD133^+^/SOX2^+^ cells and increases the CD44^+^/Chi3l1^+^ cells. Chi3l1 binds to CD44 and induces phosphorylation and nuclear translocation of beta-catenin, Akt and STAT3. Single cell RNA-seq and RNA velocity following incubation of GSCs with Chi3l1 show significant changes in GSC state dynamics driving GSCs towards a mesenchymal expression profile and reducing transition probabilities towards terminal cellular states. ATAC-seq reveals that Chi3l1 increases accessibility of promoters containing MAZ transcription factor footprint. Inhibition of MAZ directly regulates genes with highest expression in cellular clusters exhibiting significant cell state transitions. Finally, targeting Chi3l1 in vivo with a blocking antibody, resets the transcriptomic profile of glioblastoma and inhibits tumor growth. Our work implicates Chi3l1 as modulator of GSC cellular states and demonstrates pre-clinical efficacy of anti-Chi3l1 antibody treatment.

## Introduction

Glioblastoma is the most prevalent and aggressive primary brain tumor. Standard treatment of glioblastoma includes surgical resection followed by radiation in the vicinity of the resection cavity and administration of temozolomide (1). Even with this therapeutic approach, tumor recurrence is inevitable (2) mainly due to the presence of glioma stem cells (GSCs), which are characterized by high migratory potential (3), resistance to chemotherapy and radiation and the ability to form recurrent tumors (4). GSCs exhibit remarkable plasticity and dynamically transition between more differentiated and less differentiated states via mechanisms akin to cellular reprogramming (5). These mechanisms of GSC plasticity have led to examination of factors that might influence tumor propagating potential and tumor recurrence. One such concept relates to the role of tumor microenvironment to induce cellular reprogramming that results in generation of more stem-like cancer cells with the ability to propagate malignancy (6). Although each glioblastoma contains cells in multiple states, plasticity between states and the potential of a single cell to generate multiple transcriptomic and phenotypic states has been demonstrated (7). From all glioma cell phenotypes, the mesenchymal is associated with worst clinical outcomes and the main phenotypic markers expressed in mesenchymal GSCs are CD44 and Chi3l1 (7). Chitinase 3-like protein 1 (Chi3l1), also known as YKL40, is a member of an evolutionary conserved 18-glycosyl-hydrolase protein family that was originally discovered in murine mammary adenocarcinoma cells as a secretory protein (8). Chi3l1 is secreted from a variety of cells including macrophages, neutrophils, epithelial, endothelial, synovial cells as well as cancer cells. Studies have demonstrated that circulating levels of Chi3l1 are increased in many malignancies, including prostate, colon, ovary, kidney, breast, glioblastoma, malignant melanoma, and lung cancer. High levels of Chi3l1 frequently associate with poor prognosis and mortality in these cancers (9-13). In glioblastoma, Chi3l1 is a key marker for identification of the mesenchymal subtype and has been implicated its increased expression in oncogenic pathways involving the NF-κB RelB and STAT-3/RTVP-1 signaling systems that are known to drive mesenchymal differentiation (14,15).

Despite these findings, the role of Chi3l1 as paracrine modulator of GSC states and the transcriptional regulatory network induced by Chi3l1 in GSCs remain unknown. Here, using single cell RNA-seq (scRNA-seq) and RNA velocity analysis, we show that incubation of patient-derived GSCs with recombinant Chi3l1 induces significant transitions of cells towards clusters with mesenchymal transcriptomic profiles. Chi3l1 significantly reduces the probability of GSCs to transition towards terminal transcriptomic states. This suggests that a function of Chi3l1 is to modulate cellular plasticity and to restrict cell commitment. ATAC-seq of Chi3l1-treated GSCs reveals targeted chromatin remodeling, increased accessibility on promoters enriched for the Myc-associated Zinc finger protein (MAZ) motif and an increase in MAZ transcription factor (TF) activity/footprinting. Knockdown of MAZ in GSCs results in inhibition of Chi3l1-induced genes that exhibit the highest expression in GSC cellular clusters undergoing cell state transitions. *In vivo* sustained local treatment of human glioblastoma xenografts with a blocking anti-Chi3l1 antibody results in inhibition of glioblastoma growth by more than 60%. Finally, human glioblastoma xenografts treated with anti-Chi3l1 show significant reduction of the mesenchymal transcriptomic signature. Our work implicates Chi3l1 as paracrine modulator of GSC cellular states and demonstrates pre-clinical efficacy of our anti-Chi3l1 antibody to reduce glioblastoma tumor burden.

## Results

### Chi3l1 induces expression of mesenchymal markers in proneural GSCs

GSCs from patients with IDH wild type primary glioblastoma have been characterized by RNA-seq and shown to recapitulate the patient’s tumor in orthotopic xenografts (16,17). GSC1 and GSC2 both exhibit chromosome 10 deletion and amplification of chromosome 7. In addition, both GSCs show EGFR amplification, while GSC2 presents significant amplification of MDM2 (Supplementary Fig. S1A).

To determine whether the presence of Chi3l1 in the microenvironment of GSCs induces phenotypic alterations, we cultured GSCs in the presence or absence of recombinant Chi3l1 for 7 days and performed Flow Cytometry analysis for the expression of markers that define the mesenchymal and proneural phenotype (18). We show that incubation of GSCs with Chi3l1 induces significant increase in CD44+/ Chi3l1+ and Chi3l1+ expression and concomitant significant decrease in the population of GSCs expressing SOX2+/CD133+ (Fig. 1A). The significant increase in CD44 expression following incubation of GSCs with Chi3l1 was also confirmed by Western blot (Fig. 1B). To evaluate bulk transcriptomic changes following exposure of GSCs to Chi3l1, we performed differential gene expression analysis using the NanoString PanCancer Progression Panel. This returned 53 genes with significantly increased expression and 12 genes with significantly decreased expression in the Chi3l1-treated set (Supplementary Fig. S1B). To determine whether these upregulated genes were correlated with a mesenchymal phenotype, we used the TCGA RNA-Seq data for cross-validation and found significant correlation with mesenchymal gene enrichment (Supplementary Fig. S1C). Since the mesenchymal phenotype has been linked to higher “stemness” potential, we examined whether culturing of GSCs in the presence of Chi3l1 affects the ability of GSCs to self-renew. We performed *in vitro* limiting dilution analysis (19) for 14 days with and without the addition of Chi3l1 and found that Chi3l1 significantly increases (p<6e-18) the frequency of self-renewal of GSCs (Supplementary Fig. S1D).

**Figure 1.**
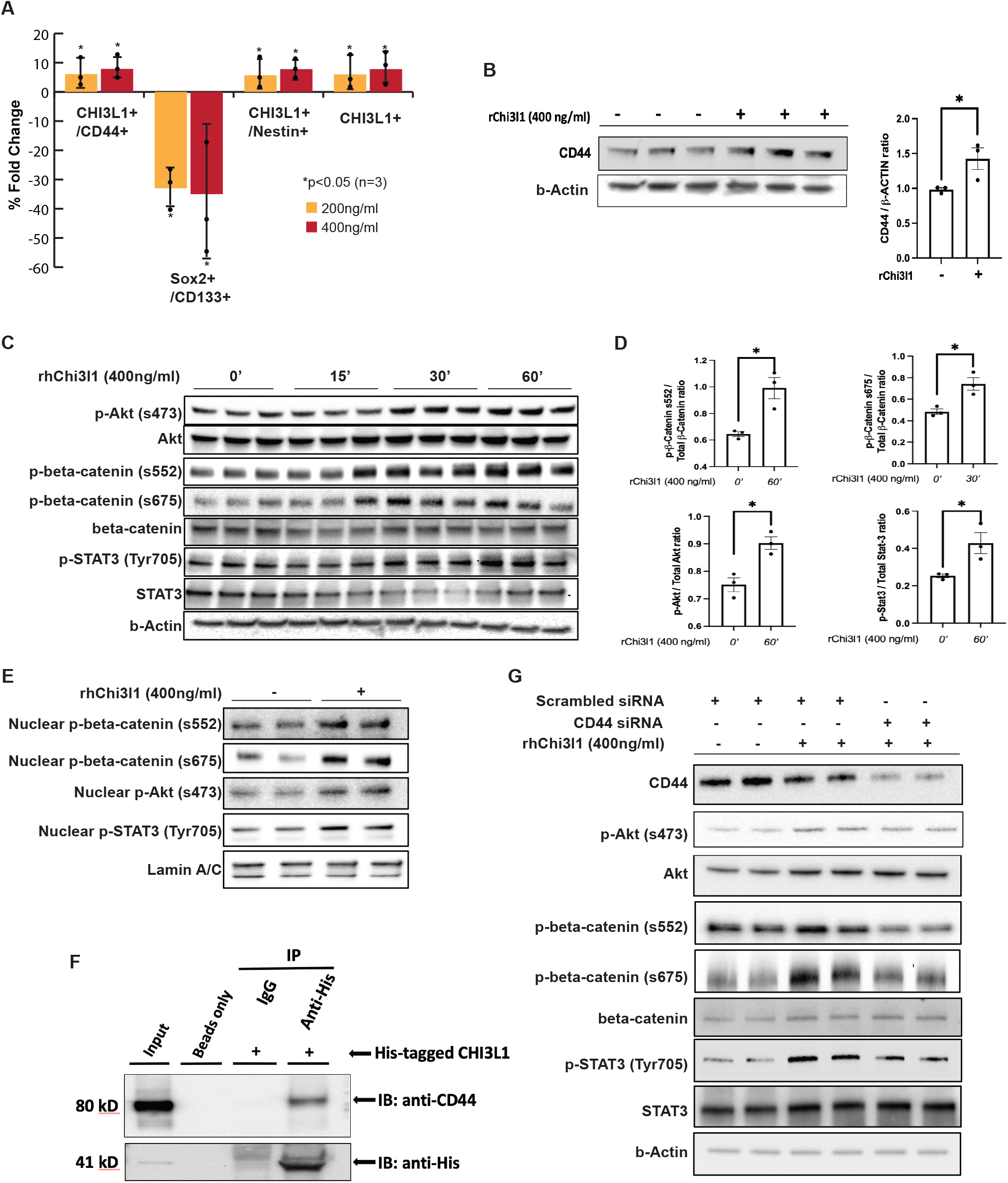
Chi3l1 interacts with CD44 and induces phosphorylation and nuclear translocation of Akt, beta-catenin and STAT3. **A**, Percentage of fold change of mesenchymal and proneural markers in GSCs following Chi3l1 treatment quantified by Flow cytometry. Student’s t-test: p-value: *<0.05, n=4. **B**, Left: Western blot analysis of CD44 in GSCs treated with or without Chi3l1. Actin was used as a loading control. Right: Densitometric analysis of the western blot results showing that GSCs treated with ChiL3 exhibit significant higher expression of CD44. The results are normalized to the expression of actin and presented as mean +/-SD from three independent experiments. Significance was calculated with Student’s t-test. **C and E**, Western blot analysis showing that incubation of GSCs with Chi3l1 induces significant phosphorylation and nuclear translocation of Akt (s473), beta-catenin (s552 & s675) and STAT3 (Tyr705). Actin (C) or Lamin A/C (E) were used as a loading control. **D**, Densitometry quantification of phosphorylation of Akt (s473), beta-catenin (s552 & s675) and STAT3 (Tyr705). Results are presented as mean +/-SD from three independent experiments. **F**, Protein lysates of GSCs (Input) were immunoprecipitated with beads only, His-tagged antibody and isotype matched IgG. Western blot was performed using CD44 antibody, which showed that Chi3l1 interacts directly and co-precipitates with CD44. **G**, Western blot analysis of phosphorylation of Akt(s473), beta-catenin(s552 & s675) and STAT3(Tyr705) in GSCs with small interfering RNA (siRNA) targeting negative control (scrambled siRNA) or CD44 and treated with or without Chi3l1 showing that knockdown of CD44 inhibits the Chi3l1 induced phosphorylation of Akt, beta-catenin and STAT3.

### Chi3l1 interacts with CD44 and induces phosphorylation and nuclear translocation of Akt, β-catenin and STAT3

In glioblastoma cells, Chi3l1 has been shown to interact with IL-13Ra2 and induce Erk1/2 and Akt phosphorylation (20). We show that incubation of GSCs with Chi3l1 induces significant phosphorylation and nuclear translocation of Akt (s473), β-catenin (s552 & s675) and STAT3 (Tyr705) (Fig. 1C, D and E). Chi3l1 interacts directly and co-precipitates with CD44 in GSCs (Fig. 1F), while direct binding and co-immunoprecipitation of Chi3l1 or CD44 with IL-13Ra2 was not detected (data not shown). Knockdown of CD44 with siRNAs inhibits the Chi3l1 induced phosphorylation of Akt, β-catenin and STAT3 suggesting that CD44 is the receptor that mediates this Chi3l1 induced signaling cascade in GSCs (Fig. 1G).

### GSC single-cell characterization

To determine the effect of Chi3l1 on GCSs at the single cell level, we applied single cell RNA-seq (scRNA-seq) on GSC1 and GSC2 treated with or without Chi3l1 for 7 days using the 10x Genomics Chromium platform. Following data integration, we sought to identify stable clusters using the Jaccard similarity index pipeline (21). Jaccard indexing is a resampling method in which 80% of cells are re-sampled from the population and re-clustered. For each cluster, a Jaccard index is calculated to evaluate similarity before and after re-clustering. Clusters with a mean stability score greater than 0.85 are highly stable while clusters lower than 0.6 are considered unstable (21). For GSC1 using k.param of 12, resolution of 0.6 and PC of 50 provides 95% of all cells within 15 highly stable clusters (Fig. 2A, B and C). For GSC2, the optimal parameters were k.param of 50, resolution of 0.6 and PC of 50, yielding in 7 stable clusters (Fig. 2D and E). Of note, cluster 6 of GSC2 was excluded from downstream analysis as it had <40 cells combined across both conditions.

**Figure 2.**
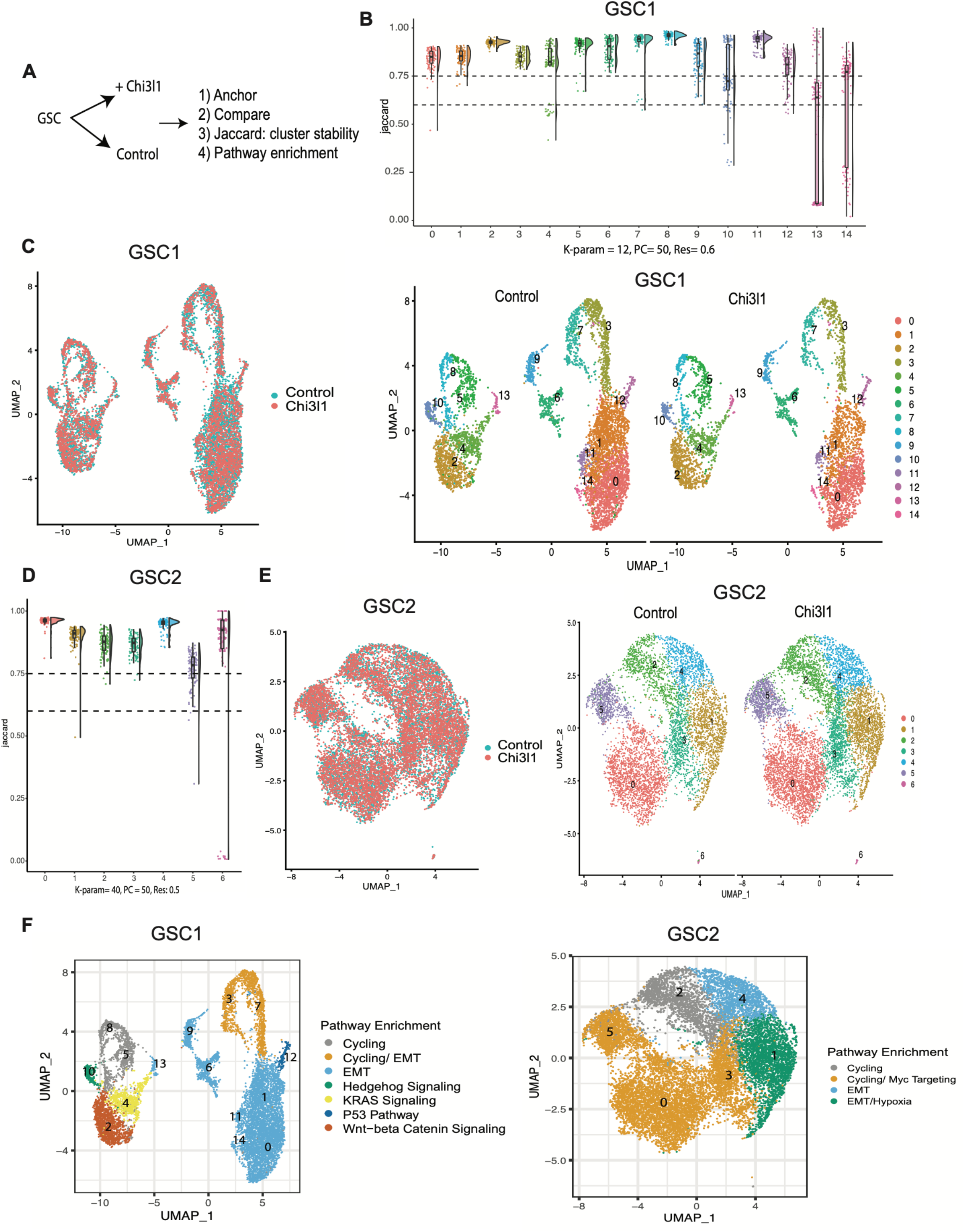
Single-cell RNA sequencing characterization of control and Chi3l1 treated GSCs. **A**, Scheme depicting the analysis set up. Briefly, GSCs incubated with or without Chi3l1 were profiled with single-cell RNA sequencing (scRNA-seq). Data from both conditions were first anchored to allow direct comparison. Second, Jaccard analysis was performed to determine stable clusters. Finally, pathway enrichment analysis was conducted per cluster. **B and D**, Jaccard Raincloud plots show the stability of each cluster for GSC1 (B) and GSC2 (D). The boxplots with a half-size violin plot displays the distribution of the Jaccard indices before and after re-clustering across 100 subsamples. The dotted lines denote Jaccard indices cutoffs at 0.6 and 0.75 (a cluster is deemed stable when its mean Jaccard index is >0.80 and unstable <0.60). **C and E**, Uniform manifold approximation and projection (UMAP) of anchored GSCs colored by condition (Left: blue = control, red = Chi3l1) and of GSCs colored by annotated stable clusters for each condition (Right). **F**, UMAP of GSCs colored by most significantly enriched pathways for GSC1 (Left) and GSC2 (Right).

Next, we performed functional enrichment analysis to characterize each cluster. GSC1 shows enrichment for six major pathways including: EMT (clusters 0, 1, 6, 9, 11, 13, 14, 7 and 3), Hedgehog signaling (cluster 10), KRAS signaling (clusters 4), p53 pathway (cluster 12), cell cycling (clusters 3, 5, 7, 8) and WNT-beta catenin signaling (cluster 2) (Fig. 2F). Notably, clusters 10, 2 and 4 express OLIG1, CCND2, PTN, GAP43, and TMSB4X. Clusters 6, 14, 1, 0 and 11 show higher expression of CD44, TGFB1, COL1A2, TFGBR2, TIMP3, SERPINF1, MGP, LGALS1 and S100A6, which are considered mesenchymal markers. Clusters 3, 8, 5 and 7 display expressions of CCNB1, PLK1, CENPE, PRC1, ASPM and CDKN3. Finally, cluster 13 exhibits a unique profile with expression of IGFBP3, SERPINE1, HLA-DRB5, SPP1, SCG2 and TGFBI (Fig. 3A). GSC2 is predominantly enriched for cell cycling (clusters 2, 5, 0 and 3) with high expression of genes such as FOXM1, followed by MYC targeting (clusters 0, 3 and 5), and EMT pathways (clusters 4 and 1) (Fig. 2F). Interestingly, clusters 2, 3, 0 and 5 are also enriched for SOX2. Cluster 4 highly expresses genes such as ANXA1, CD44, LGALS1 and CAV1, while cluster 1 expresses WISP1 and the astro-mesenchymal marker S100B (Fig. 3C). In summary, we note vastly different transcriptomic states and cellular composition between both sample (Fig. 3B and D). Ultimately, the single cell characterization of these two GSCs highlights the striking heterogeneity inherent to glioblastoma.

**Figure 3.**
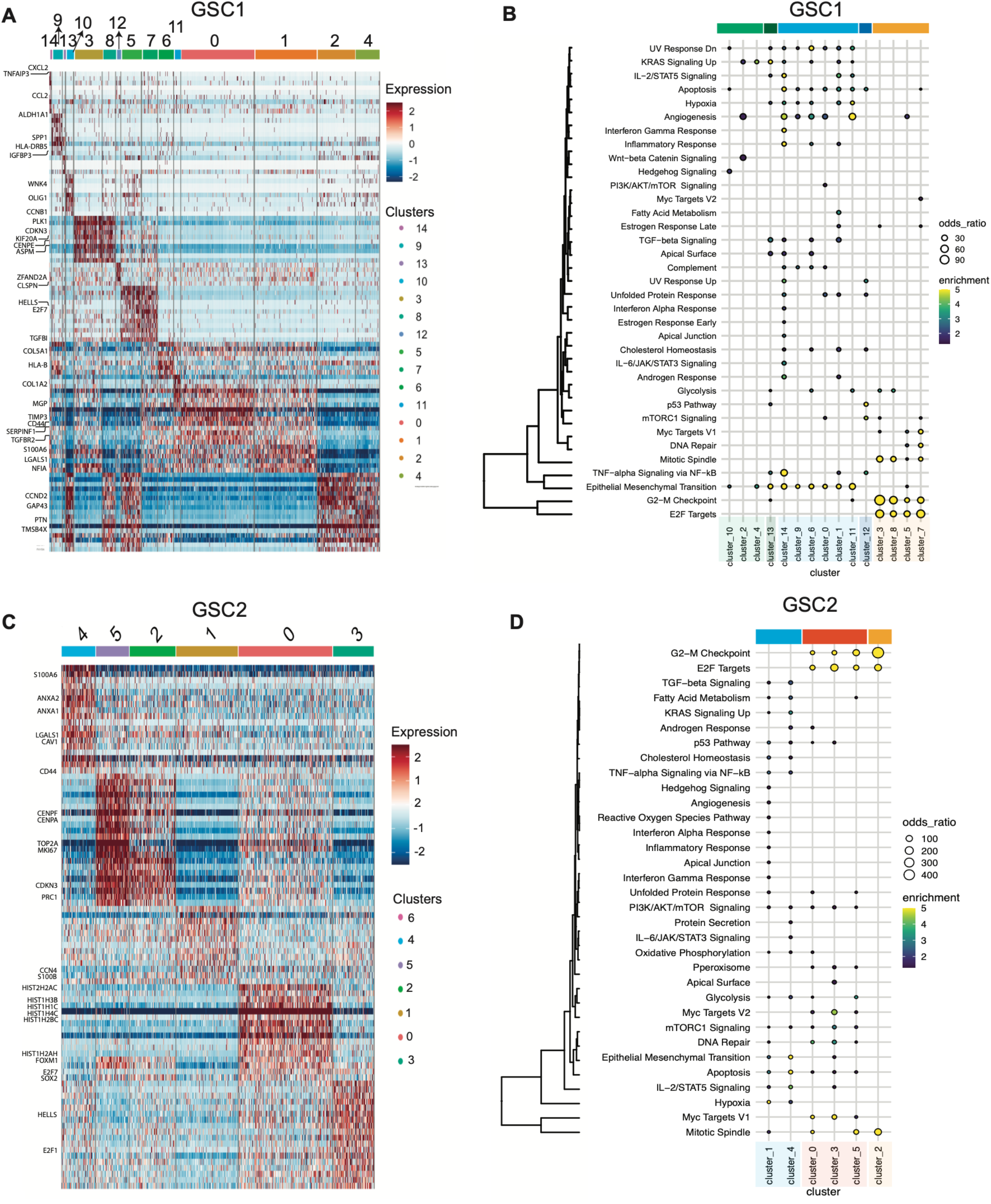
Functional enrichment analysis of scRNA-seq clusters displays the heterogeneity inherent to glioblastoma. **A and C**, Characterization heatmaps showing cluster specific gene expression profiles. **B and D**, MSigDB cancer hallmark pathway enrichment analysis. Dot plots show the most significant MSigDB pathways enriched per cluster for GSC1 (B) and GSC2 (D). The size of the dot is based on the odds ratio, which is the radio of the cluster specific, differentially expressed genes in the reference to the total number of genes in a specific pathway gene set, while the dot color gradient corresponds to the pathway enrichment, which is derived from log transformed p-values.

### RNA velocity predicts significant cell state transition changes following Chi3l1 exposure

To investigate the role of Chi3l1 on the transcriptomic state of GSCs, we employed RNA velocity analysis (the change in mRNA abundance) in order to predict cell state trajectories. To accomplish this, we used a likelihood-based dynamical model generalized to transient transcriptomic cell states. RNA velocity fields show velocity trends specific to each patient derived GSC. In GSC1, we observe pronounced RNA velocity field changes following Chi3l1 exposure, suggesting treatment induced cell state transition changes. There are striking directionality changes in RNA velocities in the cycling cluster 5 and KRAS/ EMT/IGFBP3 enriched cluster 13. Cluster 13 transitions away from KRAS enriched cluster 4 and towards EMT enriched populations. Additionally, we notice transition changes in the KRAS enriched population (cluster 4), the WNT-beta catenin (cluster 2) and EMT enriched population (cluster 9 and 6) (Fig. 4A). In GSC2, prior to Chi3l1 exposure we observe a general state transition from the initial cluster 2 towards cluster 5. However, after exposure of GSC2 to Chi3l1, the velocity field becomes less linear/direct, suggesting a disruption of the GSC2 cell-cell state transition. We note a striking increase in transition towards EMT populations, cluster 4 and cluster 1 (Fig. 4B).

**Figure 4.**
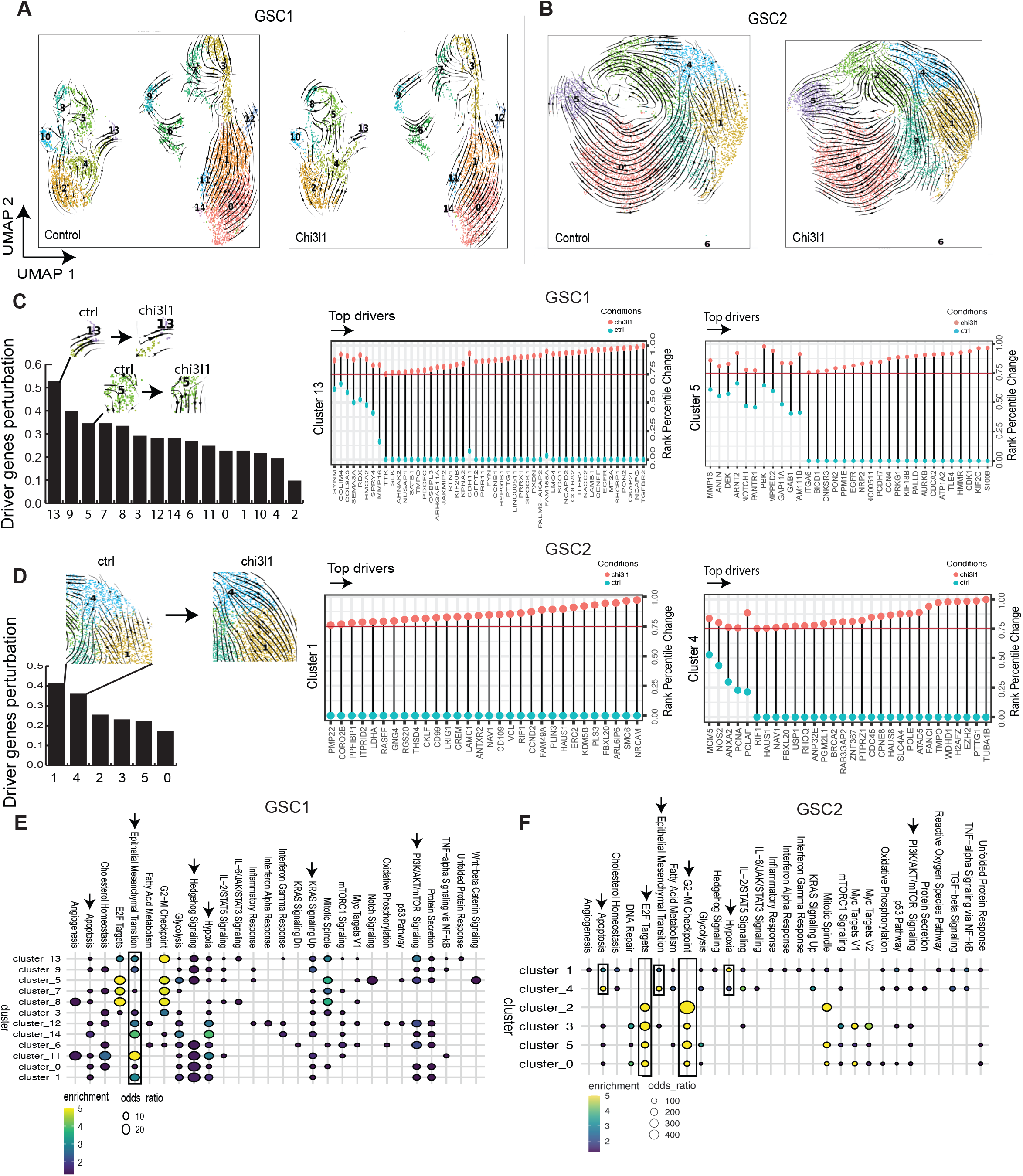
RNA velocity predicts significant cell state transition changes following Chi3l1 exposure. **A and B**, Velocity fields derived from the dynamical model (scVelo) are visualized as streamlines and projected unto UMAP embedding of anchored cells. Cells are colored by pre-defined clusters. **C and D**, Detection of driver genes. Genes were ranked based on likelihood scores and assigned a percentile rank. Perturbation percent/fraction refers to the fraction of the top quartile driver genes that are displaced as top likelihood ranking genes following Chi3l1 exposure. Bar plots display for clusters that experience the highest perturbations as well as the percentile rank of driver genes in control (blue) and after Chi3l1 exposure (red). The driver genes are ordered from left to right by higher significance (black arrow). The red line corresponds to a cut-off of 0.75 percentile rank in Chi3l1 sample. **E and F**, Genes with a percentile rank in Chi3l11 >0.75 and with a difference percentile rank between Chi3l1 and control >0.25 were considered as significant driver genes and queried using EnrichR (MSigDB) to determine pathways enrichment. Dot plots depict the MSigDB pathways enriched of these driver genes. GSC1 shows an enrichment for EMT, hedgehog signaling, KRAS, hypoxia and mTOR signaling, while GSC2 shows enrichment for cycling, hypoxia, EMT, MYC targeting and apoptosis.

To further supplement these observations, we developed an analytical pipeline (Supplementary Methods) for unbiased statistical comparison of cell transition changes across conditions. The first approach is a rank-based method, while the second employs a cluster to cluster pairwise transition probability distribution comparison. These methods were developed to allow for 1) the quantification of significant cell state transition changes across conditions and 2) to identify gene dynamics that drive these state changes.

### Chi3l1 induced state transition dynamics are driven by changes in genes enriched for EMT and stemness pathways

To evaluate our rank-based analysis, we assessed the concordance between observed velocity field changes and the percentage of top quartile genes that have been perturbed after Chi3l1 referred as percent driver gene perturbation. We establish that this percentage was consistent with velocity map changes. For example, in GSC1, cluster 13 and 5 which experience the most significant vector changes exhibit the highest percent perturbation. This is observed in cluster 1 and 4 for GSC2 as well (Fig. 4C and D). Next, we performed a pathways enrichment analysis per cluster on the genes with the most significant rank changes. These are genes that undergo a shift from having no predicted role in state transitions to being top state transition defining genes following incubation with Chi3l1. We find that the observed cell state transition changes following Chi3l1 exposure are driven by genes involved in EMT and stemness in both GSC. GSC1 shows an enrichment for EMT, hedgehog signaling, KRAS, hypoxia and mTOR signaling. GSC2 shows enrichment for cycling, hypoxia, EMT, MYC targeting and apoptosis (Fig. 4E and F).

### Chi3l1 increases transition towards EMT enriched glioma stem cell states

In GSC1, there is a notable increase in GSC state transition towards EMT enriched clusters, particularly towards cluster 1 and cluster 3. These clusters exhibit the highest scores for significant increase in incoming transitions following Chi3l1 (about 40% of all transitions are significant). Together, CD44/EMT enriched clusters 1 and 3 account for more than 1/3 of all significant increase in incoming transition (11 of 28 total significant increase in transitions in Chi3l1 relative to control) (Fig. 5A). Furthermore, we also note an equal, marked increase in transition towards EMT enriched cluster 7, cycling cluster 5 and hedgehog enriched cluster 10 (Fig. 5A and C). For GSC2, we also report an increase in transition towards the EMT enriched cluster 4. Additionally, a notable transition towards cycling/MYC enriched cluster 3 is observed (Fig. 5B and D). Taken together, the data suggest that Chi3l1 induces an increase in transition towards EMT and canonical stemness (hedgehog, MYC) enriched GSC states.

**Figure 5.**
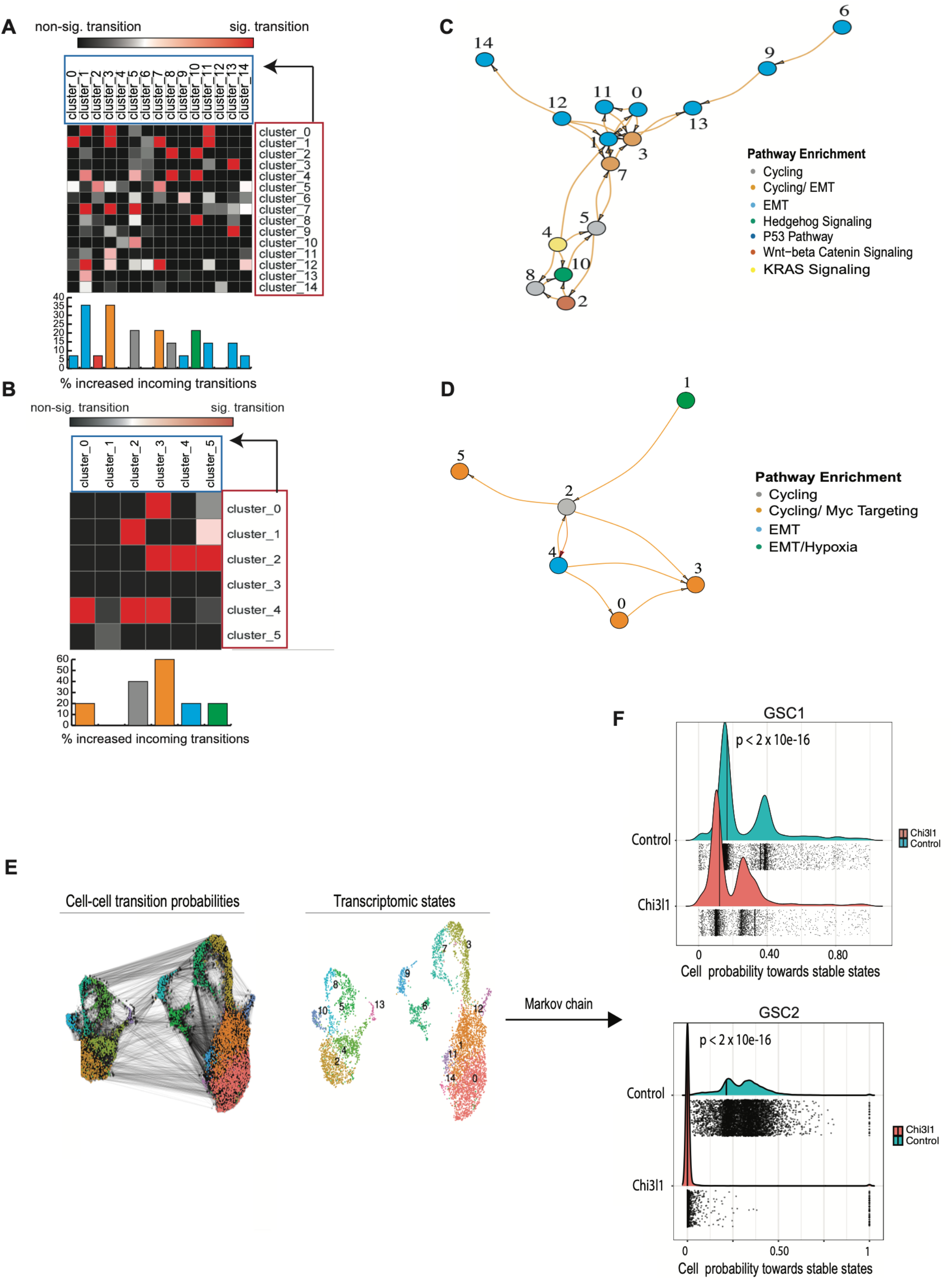
Chi3l1 increases transition towards EMT enriched GSC states and decreases transition towards stable states. **A and B**, Matrices of pairwise cluster to cluster transition probability changes (here is only shown cluster to cluster transitions that experience an increase in transition in Chi3l1 condition compared to control). Arrow denotes transition order from an initial clusters/state i (in crimson) to final state j (turquoise). The increase in transition that are statistically significant are colored in the red/pink scale, while the non-significant ones are in the black/gray scale (Pval<0.05, Mann-Whitney U Test). **C and D**, Road map visualization of significant cluster to cluster increase in transitions following Chi3l1 treatment as a nodes and edges network. **E**, Cell-cell transition probabilities were computed by integrating velocity data and cellular transcriptomic similarities. This generated general cell-fate trajectory hypotheses as displayed in the cell-cell probabilities plot. **F**, Plot shows significantly lower probabilities towards terminal states following Chi3l1 exposure for GSC1 (top) and GSC2 (bottom).

### Chi3l1 decreases transition towards stable states

RNA velocity data suggest that Chi3l1 exposure induces significant changes in GSCs state transition. To better characterize the role of Chi3l1 in cell state dynamics and supplement our previous findings, we applied a Markov-based transition probability model (CellRank) (https://www.biorxiv.org/content/10.1101/2020.10.19.345983v2.full) to our RNA velocity data. This trajectory inference model uses RNA velocity to identify “initial”/ “terminal” transcriptomic states and assigns each cell a fate probability of assuming a “terminal” transcriptomic state (Fig. 5E). Initial states refer to cell states that are predicted to be the founding or precursor populations and therefore have the lowest incoming transition probabilities. Conversely, terminal states which can be thought of as “attractor” states refer to states with highest self-transition probabilities, in other words states that are most readily assumed by the cells (22). First, we identify initial and terminal transcriptomic states (Supplementary Fig. S2A). Next, we examined transition probabilities towards terminal transcriptomic states. We find significantly lower probabilities towards terminal states following Chi3l1 exposure (Fig. 5F) for both patients with the effect even more pronounced for GSC2, suggesting a general loss of GSC commitment towards these attractor/terminal transcriptomic cellular states. In conclusion, these data suggest that Chi3l1 induces GSCs to acquire a poised, more plastic/transitional phenotype.

### Effects of Chi3l1 on chromatin accessibility and transcription factor footprinting of GSCs

To determine transcriptional regulatory mechanisms modulating the cell state transition changes following exposure to Chi3l1, we performed ATAC-Seq on Chi3l1-treated and control GSCs. Annotation of the peaks reveals that the peak densities at promoter regions increase in signal for the Chi3l1 treated group, indicating increased accessibility (Supplementary Fig. S3A). We confirmed that differential analysis was not biased towards genomic regions with intrinsically high counts by studying the differentially accessible sites relative to their average log normalized counts (Supplementary Fig. S3B). Analysis of chromatin accessibility identifies 1034 differentially accessible promoters, 525 of which with increase accessibility following Chi3l1 treatment (Fig. 6A). Motif enrichment analysis in promoters with increase in accessibility shows enrichment for SOX9, S0×10, MYB, ZNF281, WT1, CTCFL and MAZ displaying the most pronounced enrichment (Fig. 6A). Then, transcription factors (TFs) activity change was probed via DAStk, HINT-ATAC and TOBIAS. TF activity using change in motif displacement scores of 768 motifs from DAStk reveals 10 TFs with significant increase in activity after exposure to Chi3l1 (Fig. 6B). Of these TFs, MAZ has the greatest significance in terms of expression and activity (Fig. 6B, Supplementary Fig. S3C and S3D). Furthermore, TF footprinting analysis using HINT-ATAC and TOBIAS independently validates MAZ as one of the most significantly activated TFs following Chi3l1 exposure (Fig. 6D, E and F). Consensus analysis across all three platforms shows that 25% of activated TFs are called by at least two of the programs, but 75% are non-reproducible/unique to each platform. Finally, only 3 TFs are reproducibly identified across all platforms to experience a significant increase in activity following treatment: KLF15, VEZF1 and MAZ (Fig. 6C). In GSCs, KLF15 has very low expression, VEZF1 is moderately expressed and MAZ is highly expressed (Supplementary Fig. S3C).

**Figure 6.**
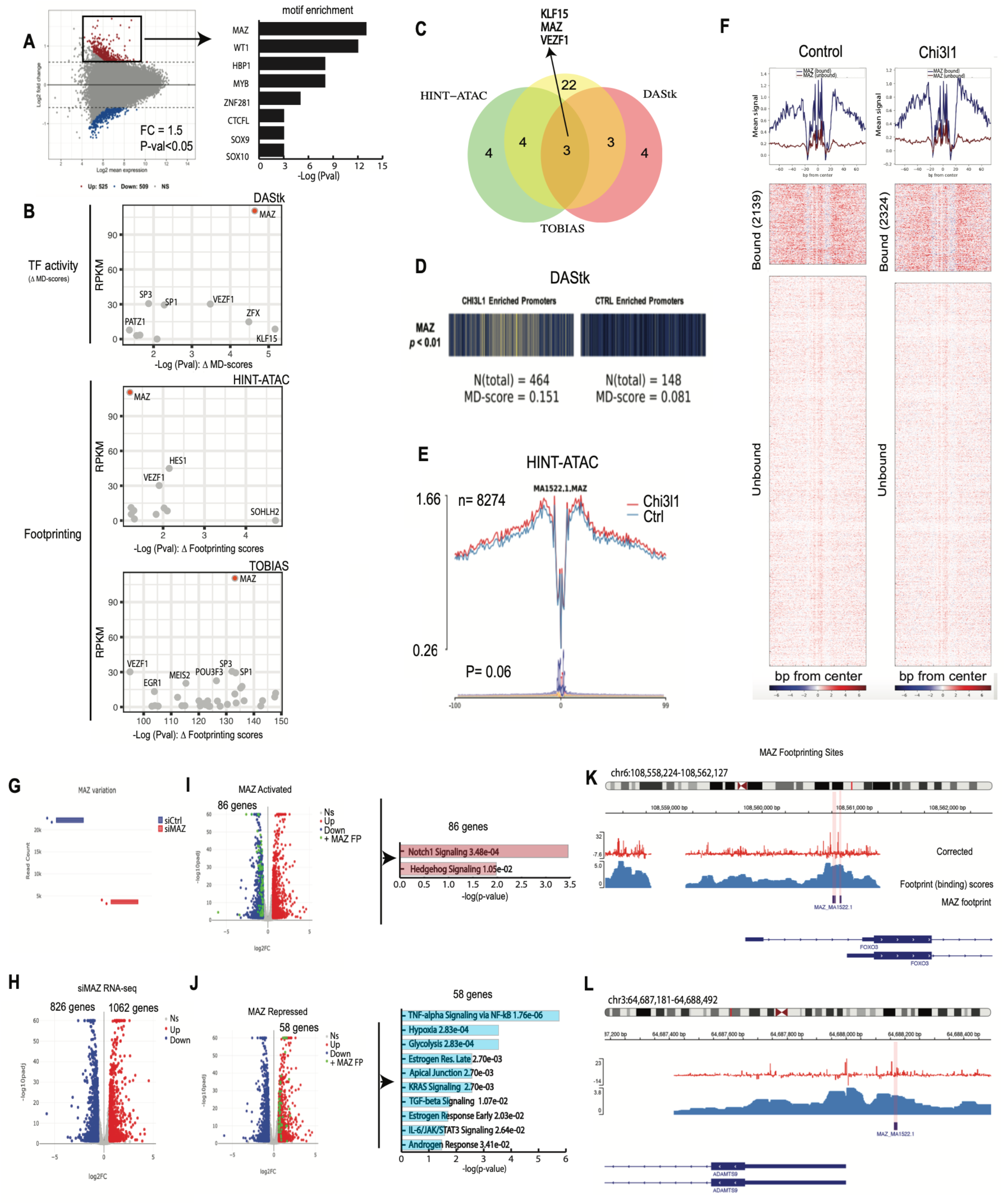
Transcription factor (TF) activity and RNA-seq integration identifies Chi3l1 activated MAZ as regulator of stemness in GSCs. **A**, MA plot exhibiting differentially accessible sites following Chi3l1 exposure (Left). The corresponding motif enrichment at promoters with accessibility gain is depicted on the right. **B**, Scatterplot of highly activated TFs expression and respective activity change following Chi3l1 exposure. TFs activity change was computed and validated using three independent TF activity platforms (upper: DAStk method with TF activity based on motif displacement or MD-scores; lower: TF footprinting based analyses (HINT_ATAC and TOBIAS differential footprinting scores)). **C**, Venn diagrams depicting TF activity consensus across the three TF activity platforms. **D**, Barcode plot encoding TF activity enrichment at MAZ motifs in Chi3l1 treated samples versus controls (Pval= 2.3E-05) using DAStk. **E**, HINT-ATAC aggregates footprint signal at MAZ motif in control versus treated samples (p=0.06)). **F**, Upper plots show aggregate signals of transcription factor binding across conditions (left: control, right: Chi3l1). Heatmaps depicted individual binding site signals show greater signal enrichment for bound sites compared to non-bound sites. Chi3l1 sample experiences an increase in bound site relative to control. **G**, Read count plot showing that MAZ is knocked down in siMAZ (siRNA). **H**, Volcano plot displays differential gene expression following MAZ knockdown (NS = non significant, up = upregulated in siMAZ, down = downregulated in siMAZ). **I and J**, show MAZ regulated genes. (Left)Volcano plot depicts differentially expressed genes following siMAZ (genes activated by MAZ (I) or repressed by MAZ (J)). Points in green emphasize differentially expressed genes with MAZ footprint/binding within their promoters. (Right)The genes were queried for pathway enrichment. **K and L**, Examples depicting footprints within the promoters of MAZ activated FOXO3 and MAZ repressed ADAMS9. Footprints coincide with sites with 1) high binding scores and 2) loss of signal within highly accessible sites that is characteristic of ATAC footprints.

### Chi3l1 induces a MAZ-regulated transcriptional network

To identify genes that are potentially directly regulated by MAZ following Chi3l1 treatment, we combined MAZ footprinting prediction from ATAC-seq data with RNA-seq data following MAZ knockdown. GSCs were treated with Chi3l1 and transfected with siCTR (n=2) and siMAZ (n=2). Following MAZ knockdown, 826 genes are downregulated and 1062 are upregulated (Fig. 6G and H). To extricate potential direct MAZ regulated genes, the promoters of differentially expressed genes following MAZ perturbation were surveyed for the presence of a MAZ footprint. We identify 3130 footprint sites or predicted binding sites. These footprints are predicted to be within the promoters of 1836 genes (representative MAZ peaks at Fig. 6K and L). Of these 1836 genes, 145 genes are found to be differentially expressed following MAZ knockdown: 86 of which downregulated so deem to be MAZ activated genes and 59 of which upregulated representing the potentially MAZ repressed genes (Fig. 6I and J). Ultimately, pathway enrichment analysis reveals that potential MAZ activated gene network is enriched for stemness maintenance mechanisms (Notch1 and Hedgehog signaling) while MAZ repressed gene network is enriched for TNF-a, hypoxia and TGF-beta signaling (Fig. 6I and J).

### MAZ regulated genes belong to clusters with cell transition changes following Chi3l1 treatment

In GSC1, approximately 60% of MAZ activated genes are relatively more expressed in populations that have experienced significant cell state changes following Chi3l1 (clusters 10, 2, 4, 5, and 8) (Fig. 7A). Interestingly clusters 5 and 10 display the highest expression of MAZ activated genes. Cluster 5 presents one of the most significant cell state changes, showing an increase in hedgehog and Notch signaling and increase in transition towards EMT and WNT-beta catenin enriched cell states following Chi3l1 exposure. Furthermore, we note increase transition from 2, 4, and 8 towards cluster 10 following Chi3l1 (Fig. 5A and C). Cluster 10, which is hedgehog enriched, shows the highest expression of the MAZ regulated genes (Fig. 7A). In GSC2, we find the MAZ genes to be highly expressed in clusters 5, 0 and 1 (Fig. 7A), which are enriched for E2F targets, EMT and cycling related genes (Fig. 4F).

**Figure 7.**
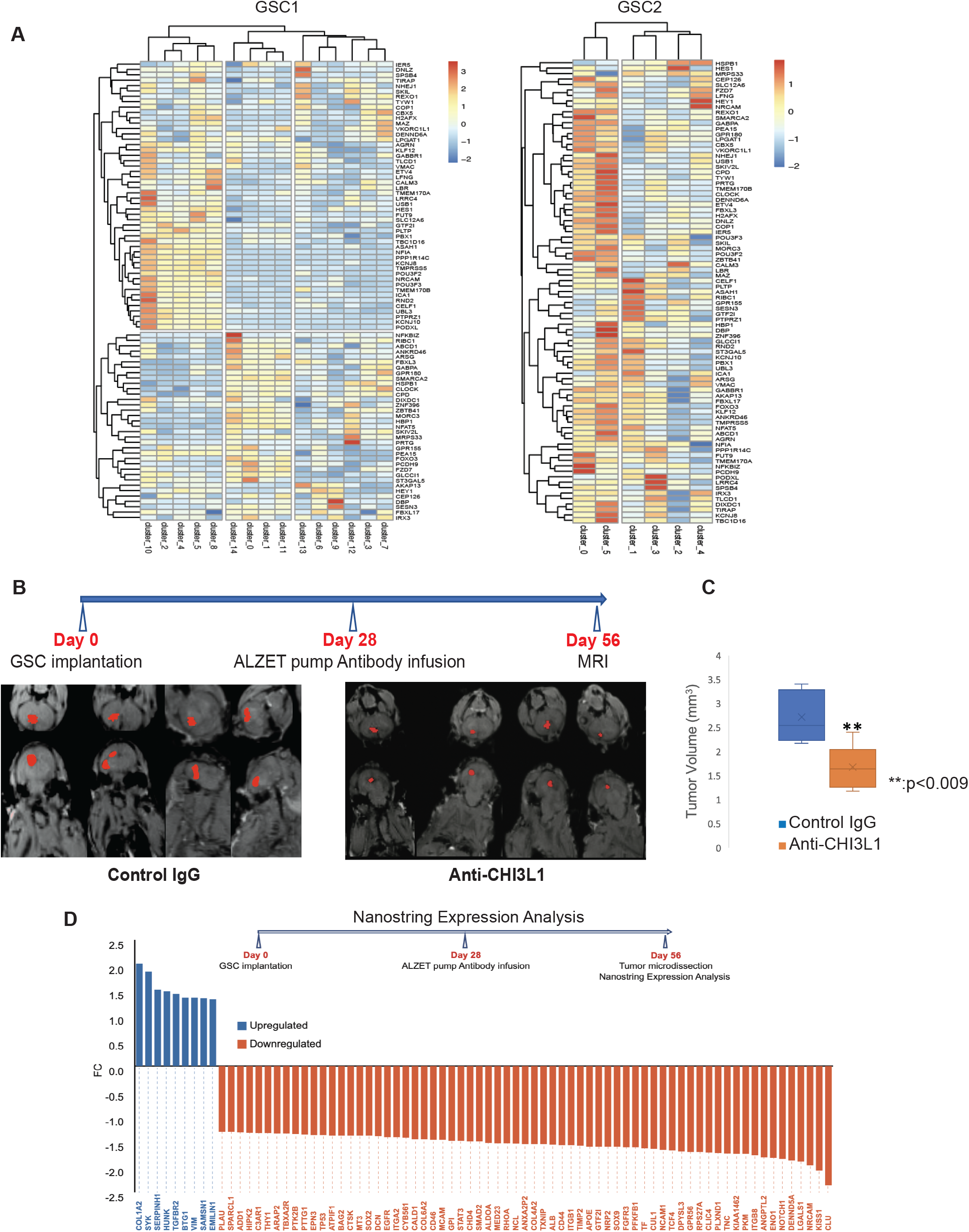
Chi3l1 induces a MAZ-regulated transcriptional network and anti-Chi3l1 blocking antibody inhibits glioblastoma growth in orthotopic xenografts. **A**, Hierarchically clustered heatmaps showing that MAZ regulated genes are relatively highly expressed in clusters that have experienced significant cell transition changes following Chi3l1 exposure. **B**, MRI images of mice treated with control IgG or anti-Chi3l1 antibody with segmented and reconstructed tumors pseudocolored in red. **C**, Volumetric surface quantification of control IgG (blue) and anti-Chi3l1 (orange) treated tumors (n=8 animals per antibody group, t-test p-value: **<0.009). **D**, Waterfall plot showing genes up- (blue) and down-regulated (orange) following anti-Chi3l1 treatment.

### Localized treatment with anti-Chi3l1 antibody results in significant reduction of human glioblastoma growth

To develop a potential therapeutic drug against Chi3l1, monoclonal blocking anti-Chi3l1 antibody isotype IgG2b, was generated and affinity purified against the peptide FRGQEDASPDRF corresponding to amino acids 223-234 of the full-length human Chi3l1 protein. Western blotting was performed to assess the specificity and sensitivity of the purified anti-Chi3l1 antibody against variable concentrations (250,125, 61.25, 31.5, 15.6, 7.8 ng/ml) of recombinant human and mouse Chi3l1 protein (Supplementary Fig. S4A). Determination of the dose response curve of the antibody displays high sensitivity in detecting Chi3l1 (Kd = 1.1⨯10^−8^) and a limit of detection of 15ng/ml (R^2^=0.991) as calculated by competitive ELISA (Supplementary Fig. S4B and C). To determine the *in vivo* efficacy of anti-Chi3l1 antibody as localized treatment for glioblastoma, we generated orthotopic xenograft glioblastoma in immunocompromised mice using patient derived GSCs as shown before (16,17). 28 days following the stereotactic implantation of GSCs, mice were implanted with an intracranial catheter connected to an osmotic pump (Alzet) for continuous infusion of anti-Chi3l1 antibody. After 28 days of continuous treatment, mice were subjected to magnetic resonance imaging (MRI) to determine the effect of anti-Chi3l1 on human glioblastoma growth. Volumetric measurements and quantification of tumor surface of mice treated with isotype matched IgG (n=8) was compared to mice treated with anti-Chi3l1 antibody (n=8) (Fig. 7B) using 3D Slicer (v.4.10.2, Brigham & Women’s Hospital). We show that continuous localized treatment with anti-Chi3l1 antibody results in more than 60% reduction of tumor volume compared to IgG treated control mice (Fig. 7C, Student’s t-test *p-*value: ** <0.009).

### *In vivo* treatment with anti-Chi3l1 inhibits the Chi3l1 regulated transcriptome of glioblastoma

To determine whether the effect of anti-Chi3l1 *in vivo* is the result of specific inhibition of the Chi3l1-induced expression of mesenchymal transcripts, we treated mice carrying orthotopic patient-derived glioblastoma xenografts with anti-Chi3l1 or isotype matched IgG as described above. At the end of the 28-day continuous treatment, we micro-dissected IgG-treated (n=5) and anti-Chi3l1-treated (n=4) tumors and performed NanoString expression analysis. We show that treatment with anti-Chi3l1 significantly downregulates the expression of transcripts with major roles in mesenchymal glioblastoma such as CD44, TNC, ANXA2, ITGB1, SMAD and STAT3 as well as transcripts that have been linked to migration, invasion and stemness of glioblastoma including SOX2, SOX9, EGFR, NOTCH1, TCF4 and MED23 (Fig. 7D).

## Discussion

Functional heterogeneity in glioblastoma is defined not only by the genetic makeup of glioma cells but also through microenvironment-driven epigenetic influences that regulate glioma cell stemness. Recently, the hierarchical cancer stem cell model has been challenged by evidence suggesting that cancer stem cells may not constitute a defined cellular entity, but rather a cellular state adapting to autocrine and microenvironmental cues (6). The result of these epigenetic influences is activation of differentiation and dedifferentiation transcriptional programs, mechanisms of chromatin remodeling, non-coding RNA species and post-transcriptional RNA modification mechanisms that regulate phenotypic states of GSCs and confer tumor recurrence and therapeutic resistance.

Chi3l1, a member of the chitinase-like glycoprotein family, was first identified from the medium of a human osteosarcoma cell line MG-63 (23). Multiple studies have shown that high serum levels of Chi3l1 are correlated with metastasis and poor survival in a broad spectrum of human carcinomas including breast cancer (24), colorectal cancer (25), ovarian cancer (26), leukemia (27), lymphoma (28), and glioblastoma (29). Chi3l1, highly expressed in the microenvironment of human glioblastomas, is known to be associated with GSCs with a mesenchymal transcriptomic profile (7,30). Phenotypic plasticity in human tumors can be driven by activation of epithelial-to-mesenchymal transition (EMT) (31). This is the process by which cells acquire plasticity and gain the properties of stem cells (32,33). EMT is linked with an undifferentiated cellular state, including the capacity for extended self-renewal and the acquisition of a stem-like gene expression program. Protein-protein interaction between Chi3l1 and CD44 at the surface of GSCs, activates a p-Akt, p-beta-catenin signaling cascade, which in turn induces CD44 expression in a pro-mesenchymal feed forward loop. Using RNA velocity at a single cell resolution, we demonstrate that the presence of Chi3l1 in the environment of GSCs induces marked transcriptomic cell state transitions towards mesenchymal phenotypes. Chi3l1 is a highly potent inducer of cellular plasticity and GSCs exposed to Chi3l1 express genes that define a “poised” uncommitted cancer stem cell state.

Reversible transitions between epigenetic states enable tumor cells to adapt to different microenvironment pressures, including therapy (34,35). GSCs employ adaptive chromatin remodeling to drive cellular plasticity and therapy resistance (36). These types of chromatin alterations could also contribute to emergence of subclones in tumors with the least ability for reversion of stem-like properties or with the highest propensity to acquire stem-like characteristics. We establish here that Chi3l1 as constituent of the GSC microenvironment, induces chromatin remodeling to facilitate accessibility of regulatory elements on promoters of genes with significant roles in the pathobiology of glioblastoma. In particular, we identify increase in MAZ transcription factor footprint in accessible chromatin regulatory regions following exposure of GSCs to Chi3l1.

MAZ is a zing finger transcription factor that is associated with Myc gene expression (37) and regulates a wide variety of genes. Recently, it was shown that MAZ interacts with CTCF and functions as a genome-wide insulator arresting cohesin movement and directly interacting with Rad21 (38). Interestingly, the second most significantly induced transcription factor following Chi3l1 treatment of GSCs is Vezf1, which has been also shown capable of pausing RNA Pol II and affecting global RNA splicing (39). It is possible that Chi3l1 induces global architectural changes in the genome through alteration of MAZ transcriptional regulation and splicing de-regulation through Vezf1. Indeed, MAZ knockdown in GSCs affects genes that belong to the clusters of cells that show the most significant changes in cell transition probabilities following exposure to Chi3l1. We posit that the Chi3l1 induced MAZ footprint in GSCs results in stabilization of CTCF binding and in combination with increased Vezf1 binding could block RNA polymerase II advancement resulting in alternative RNA splicing patterns. The significant cell state transition changes noted after Chi3l1 treatment of GSC could represent global genomic effects of MAZ and Vezf1 since they reflect RNA splicing ratios inferred as RNA velocity patterns.

Elucidating state transition programs and mechanisms driving cellular plasticity will be essential to overcome current therapeutic limitations in glioblastoma. We show here that blockade of Chi3l1 using a highly specific humanized monoclonal antibody reverts pro-mesenchymal transcriptomic transitions in human glioblastoma xenografts and significantly reduces tumor burden. It remains unclear whether the microenvironment selects for survival of specific GSCs or whether tumor cells adapt within new microenvironments. We favor the later and demonstrate that Chi3l1 is a microenvironment factor that mediates cell state transitions in glioblastoma driving GSCs towards a poised uncommitted fate. We envision anti-Chi3l1 as a potent therapeutic agent in combination therapies targeting cellular adaptation in glioblastoma.

## Author Contributions

CG, BA, DK, JC, SK, EF, RS and AF performed experiments and analyzed data; CG, and DK wrote parts of the manuscript; SAT edited the manuscript, provided recourses and acquired funds; CL and JE provided resources and edited the manuscript; NT conceived the project, interpreted and analyzed data, wrote the manuscript, acquired funds, provided resources, supervision and mentoring.

## Acknowledgments

The authors would like to thank Owen Leary for analyzing the mouse MRI images with 3D Slicer. This work was supported by the National Cancer Institute R21CA235415 to N.T., by Warren Alpert Foundation Grant to N.T and S.A.T. and by funds of the Neurosurgery Department of Brown University to N.T.

## Materials and code availability

Anti-Chi3l1 neutralizing antibody (called FRG) will be made available to all investigators from academia after signing a Material Transfer Agreement (MTA) as long as the planned experiments do not directly conflict with experiments from collaborators with already established MTAs. Details for the use of code supporting data analysis of the current study are available in Method Details and from the corresponding author upon request. The data discussed in this manuscript have been deposited in NCBI’s Gene Expression Omnibus and are accessible through GEO Series accession number GSE154060 (https://www.ncbi.nlm.nih.gov/geo/query/acc.cgi?acc=GSE154060)

## Methods

### Isolation and culture of patient derived GSC cultures

The IRB of Rhode Island Hospital has approved the collection of patient-derived glioblastoma multiforme tissue. All collections were performed with patient consent and in completely de-identified manner. Primary hGSC spheres were cultured from human glioma samples as previously described (16). All hGCs used in this study were authenticated by ATCC using short tandem repeat (STR) analysis. GSCs used were between passages 5 and 30 and cultured either as spheres or as attached on fibronectin-coated plates (10ug/ml, Millipore Sigma) in a medium of 1X Neurobasal Medium, B27 serum-free supplement, minus Vitamin A, 100X Glutamax (Fisher Scientific), 1mg/ml Heparin (STEMCELL Technologies), 20 ng/ml epidermal growth factor (Peprotech), 20ng/ml basic-fibroblast growth factor (Peprotech).

### Mouse Studies

All animal experiments were approved by Rhode Island Hospital’s Institutional Animal Care and Use Committee (IACUC) and conformed to the relevant regulatory standards and overseen by the institutional review board. 9 weeks old female and male NU/J homozygous mice were obtained from Jackson Laboratory and used for the stereotactic injections. Animals were housed together until the day before the surgery and then housed individually. Animals receiving treatment were randomly assigned.

**Flow Cytometry, Western Blot, Immunoprecipitations, Nanostring expression analysis, Bulk RNA-seq analysis, Anti-Chi3l1 antibody characterization, and Methods for stereotactic implantation and MRI imaging are described at Supplementary Data**.

### ATAC-Seq

#### Sample Preparation

ATAC-seq libraries were prepared according to the protocol established and optimized before (40). Specifically, cell lysis was performed using 50ul of cold lysis buffer (resuspension buffer, 0.1% NP-40, 0.1% Tween-20, and 0.01% Digitonin (Promega)), and transposition of the resulting nuclei pellet from the lysate by Tn5 transposase was accomplished using the Nextera DNA Library Prep Kit (Illumina). DNA isolation from the transposition reaction was done using MinElute Reaction Cleanup Kit (Qiagen), and the library was generated from the isolated DNA through PCR amplification protocol. Purification of the library for optimal library preparation included removal of primer dimers and DNA fragments larger than 1000 base pairs (bp) through double-sided bead purification with Agencourt AMPure XP beads (Beckman Coulter). Sequencing of the final library was performed by GENEWIZ using the Illumina HiSeq 2500 system.

#### ATAC-Seq FASTQ Alignment, BAM Preprocessing, and Peak Calling

FASTQ files of sequencing reads for each sample were filtered, deduplicated, and aligned to the GRCh38/hg38 human reference genome using the paired-end alignment feature. BAM files from FASTQ alignment were sorted, indexed, and isolated for uniquely mapped reads using the SAMtools. Finally, peak calling was performed using Genrich with a false discovery rate (FDR) adjusted p-value significance threshold of 0.05 (-q 0.05) and a minimum peak length of 200 bp (-l 200). Consensus peak set was derived for each group by performing individual peak regions based on Fisher’s combined probability test.

#### ATAC-Seq Analysis

ATAC-Seq data analyses were performed using R. The processing of the narrowPeak files into GRanges object was done using *readPeakFile. Peaks were annotated to* “TxDb.Hsapiens.UCSC.hg38.knownGene” database and visualized with *plotAvgProf2*.

#### Accessibility changes in promoters following addition of Chi3l1

To identify differentially accessible chromatin regions, sequence read counts of each peak range in the consensus peakset for each sample were calculated with *featureCounts*. The resulting counts matrix was then converted into a DESeq data set and normalized via Median-by-Ratio Normalization (MRN) for differential chromatin accessibility analysis by DESeq2 (41). Differentially accessible chromatin regions were defined as peak regions with a normalized count fold change greater than 1.5 and a *p*-value smaller than 0.05. Differentially accessible chromatin regions were then annotated to promoters as followed: 1000 bp upstream and downstream of a TSS; the flanking radius to search beyond the promoter was 3000 bp upstream and downstream

#### Motif enrichment and transcription factor binding analysis

Motif enrichment within highly accessible promoter sites was performed using the HOMER motif analysis package with the following parameters: “hg38 -len 8,10,12 - mask -size given”. Both *de novo* and known motif enrichment results were considered. Three platforms were used for consensus differential activity analysis: the Differential ATAC-seq Toolkit (DAStk), Hmm-based IdeNtification of Transcription factor footprints (HINT-ATAC) and Transcription factor Occupancy prediction by investigation of ATAC-seq signal (TOBIAS)(42-44). Enriched motifs were probed for differential TF activity inference using DAStk differential MD score. MD scores were obtained by assessing the density of biologically active motifs active at enhancer sites (eRNA). Motif activity and hence TF binding probability was defined by the degree of motif colocalization with an enhancer RNA origin as previously described (45). To detect footprints, TOBIAS and HINT-ATAC were independently ran with respectively JASPAR CORE motif set and HOCOMOCO.

### Single-cell RNA-seq

#### Sample preparation

GSCs were treated for a week with or without Chi3l1 (500ng/ml). scRNA-seq libraries were generated using the Chromium Next GEM Single Cell 3’ GEM, Library & Gel Bead Kit v3.1, Chromium Single Cell 3’, Dual Index Kit TT Set A, Chromium Next GEM Chip G, and 10x Chromium Controller (10x Genomics) as per 10x Single Cell 3’ protocol. The sequencing was done by the Molecular Biology Core Facilities at Dana-Farber Cancer Institute using Illumina NovaSeq SP Flowcell (800M Reads) as well as the raw data processing. CellRanger count outputs were provided.

#### Analysis of scRNA sequencing data

##### Data integration and clusters determination

Using 10x Genomics platform, for GSC1/GSC2, 8000/7015 cells and 7112/8961 cells passed the sequencing threshold QC for the non-treated/control and Chi3l1 treated samples respectively. Then, low complexity cells (<1000 genes) and cells with >12% mitochondrial genomic content were excluded as previously described for GBM (46) resulting in a final population of 6102/5892 and 4060/7558 cells for control and Chi3l1 samples for GSC1/GSC2 respectively.

Seurat (47) was used for downstream analysis. First each library was converted into a Seurat object using read10x and makeseuratobject. Data was normalized using SCTransform method then objects were integrated along identified anchors to identify cell types across conditions. To determine optimal parameters for dimensional reduction and evaluate cluster stability, Jaccard similarity index was applied involving a snakemake pipeline to scatter and gather Seurat object by subsampling the cells and repeat for multiple times as described in (21) with the following parameters: subsample rate: 0.8; number of subsample: 100; subsample ks: 8, 10, 12, 14, 16, 20, 30, 50, 80, 100; subsample resolutions 0.5, 0.6, 0.8, 1, 1.2, 1.5; subsample pcs: 20, 30, 40, 50. Cells were then classified based on top expressed genes and functional pathway enrichment. First, the cluster defining genes across conditions were identified using the FindConservedMarkers. Second, the most highly expressed genes per cluster were extracted from the Seurat top ranked “identity”. Third, MSigDB cancer hallmark geneset was used in enrichr platform for cluster specific functional enrichment analysis.

##### RNA velocity and trajectory inference

RNA velocity analysis was performed using both velocyto and scVelo. First, the cellranger count output was pre-processed using the velocyto command-line tool (CLI) with default parameters specific for 10x Genomics data (48). The .loom files generated were filtered for complexity and mitochondrial content and were then loaded to scVelo (49) for downstream RNA velocity analysis. The data was preprocessed by selection the top 2000 variable genes with minimum shared counts of at least 20 and log normalized for cell size. RNA velocity was estimated using the dynamical mode and projected unto anchored Seurat derived UMAP embeddings.

CellRank was used to predict overall cell state trajectories and cell-cell transition probabilities. Genes were ranked based on likelihood scores and assigned a percentile rank. Differential ranking analysis was performed, cluster by cluster, by calculating the difference percentile rank between Chi3l1 vs control. EnrichR (MSigDB) was queried for pathways enrichment. Furthermore, a method was developed to characterize transition probabilities between cluster (Supplemental data). Generalized Perron Cluster Analysis (GPCCA) (50) was used to identify initial and terminal states as well as single cell probabilities toward terminal states.

### General statistical analysis methods

Our goal is to obtain results with greater than 95% confidence level. Assuming that data were normally distributed and that the standard deviation for measurements was no more than 3/4 of the mean, the t-test of mean was used to estimate the number of required observations. For FACS, we used n=4 biological replicates, for NanoString analysis using GSCs, we used n=3 biological replicates and GSCs from two patients. For MRI quantification we used n=8 animals per group, while for NanoString *in vivo* analysis we used n=5 IgG control treated mice and n=4 anti-Chi3l1 treated mice. Age-matched female and male mice were randomly used for *in vivo* experiments and the quantification of tumor area was performed blindly in regard to treatment group assignment by two independent researchers.

To determine significance among the means of three or more independent groups, we used one-way ANOVA. The homogeneity of variances was confirmed with Brown and Forsythe test, and the significance between specific groups was calculated with a post hoc Dunnett test. To determine significance among the means of two independent groups, we performed an unpaired two-tailed t test. To verify Gaussian distribution of data before applying the t test, we performed the D’Agostino and Pearson and Shapiro-Wilk normality tests.

